# ERECTA family signaling constrains *CLAVATA3* and *WUSCHEL* to the center of the shoot apical meristem

**DOI:** 10.1101/2020.02.24.962787

**Authors:** Liang Zhang, Daniel DeGennaro, Guangzhong Lin, Jijie Chai, Elena D. Shpak

## Abstract

The shoot apical meristem (SAM) is a reservoir of stem cells that gives rise to all post-embryonic aboveground plant organs. The size of the SAM remains stable over time due to a precise balance of stem cell replenishment versus cell incorporation into organ primordia. The WUSCHEL (WUS)/CLAVATA (CLV) negative feedback loop is central to SAM size regulation. Its correct function depends on accurate spatial expression of *WUS* and *CLV3*. A signaling pathway, consisting of ERECTA family (ERf) receptors and EPIDERMAL PATTERNING FACTOR LIKE (EPFL) ligands, restricts SAM width and promotes leaf initiation. While ERf receptors are expressed throughout the SAM, EPFL ligands are expressed in its periphery. Our genetic analysis demonstrated that ERfs and CLV3 synergistically regulate the size of the SAM, and *wus* is epistatic to *erfs*. Furthermore, activation of ERf signaling with exogenous EPFLs resulted in a rapid decrease of *CLV3* and *WUS* expression. ERf-EPFL signaling inhibits expression of *WUS* and *CLV3* in the periphery of the SAM, confining them to the center. These findings establish the molecular mechanism for stem cell positioning along the radial axis.

**Summary statement:** ERf signaling restricts the width of the shoot apical meristem, a structure which generates aboveground plant organs, by inhibiting expression of two principal regulators, *CLV3* and *WUS*, at its periphery.

## INTRODUCTION

The shoot apical meristem (SAM) generates new organs throughout the life of a plant. As stem cells in the central zone of the dome-shaped SAM slowly divide, some of their progeny are displaced laterally into the peripheral zone and basally into the rib zone. Cells in the peripheral and rib zones rapidly divide, differentiate, and are incorporated into forming leaves, flowers, and stems. Even though cells are constantly dividing, the SAM size remains stable throughout development due to a tight balance of proliferation and incorporation of cells into new organs.

The principal regulator of SAM size is a negative feedback loop consisting of WUSCHEL (WUS) and CLAVATA3 (CLV3). WUS is a homeodomain transcription factor that maintains the pool of stem cells; in its absence stems cells arise but almost immediately differentiate (Laux et al., 1996). *WUS* is expressed in the organizing center beneath the central zone, and the protein moves up into the central zone through plasmodesmata (Brand et al., 2000; Daum et al., 2014; Mayer et al., 1998; Schoof et al., 2000; Yadav et al., 2011). *CLV3* encodes a secreted peptide expressed in the central zone and perceived by multiple plasma membrane localized receptors: CLV1, CLV2, BARELY ANY MERISTEM 1 (BAM1), BAM2, CORYNE (CRN), RECEPTOR-LIKE PROTEIN KINASE 2 (RPK2), and CLAVATA3 INSENSITIVE RECEPTOR KINASEs (CIKs) (DeYoung et al., 2006; Fletcher et al., 1999; Hu et al., 2018; Kinoshita et al., 2010; Muller et al., 2008; Shinohara and Matsubayashi, 2015). In the central zone, WUS binds directly to the promoter of *CLV3* and activates its expression while CLV3-activated signaling inhibits *WUS* expression (Brand et al., 2000; Schoof et al., 2000; Yadav et al., 2011), forming a regulated feedback loop.

One of the central questions to understanding the meristem is how the spatial expressions of *CLV3* and *WUS* are established and maintained in the face of the continual turnover of cells. Previous studies have elucidated key mechanisms underlying the apical-basal distributions of *CLV3* and *WUS*. The depth of the *WUS* expression domain is defined by opposing activity of CLV3 and cytokinin: CLV3 inhibits *WUS* expression while cytokinin signaling promotes it. Both signals are produced in the apical region of the meristem and form a diffusion gradient along the apical basal axis. Cytokinin is perceived in deeper tissue layers than CLV3 which establishes *WUS* expression at a certain distance from the surface of the SAM (Chickarmane et al., 2012). Confinement of *CLV3* expression to the region above the *WUS* domain is dependent on HAM1, a transcription factor in the rib zone. Interaction with HAM1 prevents WUS from activating CLV3 transcription, which restricts CLV3 expression to the apical region of the SAM (Zhou et al., 2018). It remains, however, unclear why *WUS* and *CLV3* are expressed only around the central vertical axis of the SAM. Previous mathematical models have used implicit or explicit assumptions to define the lateral boundary that confines the expression of *WUS* and *CLV3* (Gruel et al., 2018; Zhou et al., 2018), but little is known about the actual existence of such a lateral signal. Here we present data showing that ERECTA family signaling restricts *WUS* and *CLV3* expression laterally, confining them to the center of the meristem thereby providing a key mechanism for SAM maintenance.

*ERECTA, ERECTA-LIKE 1 (ERL1)*, and *ERL2*, collectively called *ERfs*, encode plasma membrane localized leucine-rich repeat receptor-like kinases (Shpak et al., 2004). The activity of ERf receptors is regulated by a group of cysteine-rich peptides belonging to the EPIDERMAL PATTERNING FACTOR/EPF-LIKE (EPF/EPFL) family (Hara et al., 2007; Hara et al., 2009; Lee et al., 2012; Lin et al., 2017). A mitogen-activated protein kinase cascade consisting of YODA, MKK4/5/7/9, and MPK3/6 functions downstream of the receptors (Bergmann et al., 2004; Lampard et al., 2009; Lampard et al., 2014; Meng et al., 2012; Wang et al., 2007). ERf signaling controls various developmental processes including stomata formation, above-ground organ elongation, SAM size, leaf initiation, and phyllotaxy (Chen et al., 2013; Shpak, 2013; Uchida et al., 2013). In the SAM, three ERfs function redundantly with single and double mutants having no or extremely weak meristematic phenotypes. Altered meristem development can be observed in the *er erl1 erl2* mutant which has a wider vegetative SAM and forms fewer leaves at almost random divergence angles (Chen et al., 2013; Uchida et al., 2013). Recently we demonstrated that ERf activity in the SAM is controlled by four ligands: EPFL1, EPFL2, EPFL4, and EPFL6, which are expressed at the periphery of the SAM (Kosentka et al., 2019). Based on altered expression of DR5rev:GFP and PIN1pro:PIN1-GFP markers in *er erl1 erl2*, we proposed that the decrease in leaf initiation might be a result of altered auxin distribution (Chen et al., 2013), but the cause of the increased meristem size, however, has remained unknown. A decrease in organ initiation does not automatically lead to an increase of SAM size. For example, the inflorescence meristem of the *pin1* mutant does not show alteration of size or WUS expression, while failing to produce flower organs (Vernoux et al., 2000). Since regulation of SAM size depends on the CLV3/WUS feedback loop, we investigated whether ERfs genetically interact with these two genes and alter their expression. Our experiments indicate that ERfs are important modulators of *CLV3* and *WUS* expression. We propose that ERf and EPFL are a part of a new regulatory circuit that enables communication between the peripheral and the central zones and specifies the location and size of the stem cell population in the SAM.

## RESULTS

### ERf/EPFL and CLV3 signaling synergistically restrict SAM size

Comparison of *clv3* and *er erl1 erl2* mutants suggests that while both ERfs and CLV3 control SAM size, they play dominant roles during different developmental stages. At one day post germination (DPG) the SAM of *er erl1 erl2* is considerably wider (105.1 ± 3.3; mean ± SE) than in the wild type (56.1 ± 1.7; p < 1.2×10^−13^ in Student’s t-test) or in *clv3* (83.0 ± 1.4; p< 4.2×10^−7^) (Fig. 1A and C), suggesting a key role for ERfs in restricting SAM size during embryogenesis. For the first five days after germination the wild type and *er erl1 erl2* SAMs do not substantially further increase in width, while *clv3* SAM size continues to increase, indicating that post embryogenesis CLV3 signaling plays the primary role in SAM size maintenance (Fig. 1A). The pathways also contribute differently to leaf initiation, with ERfs promoting leaf initiation and CLV3 slightly inhibiting it (Fig. 1B).

**Fig. 1.**
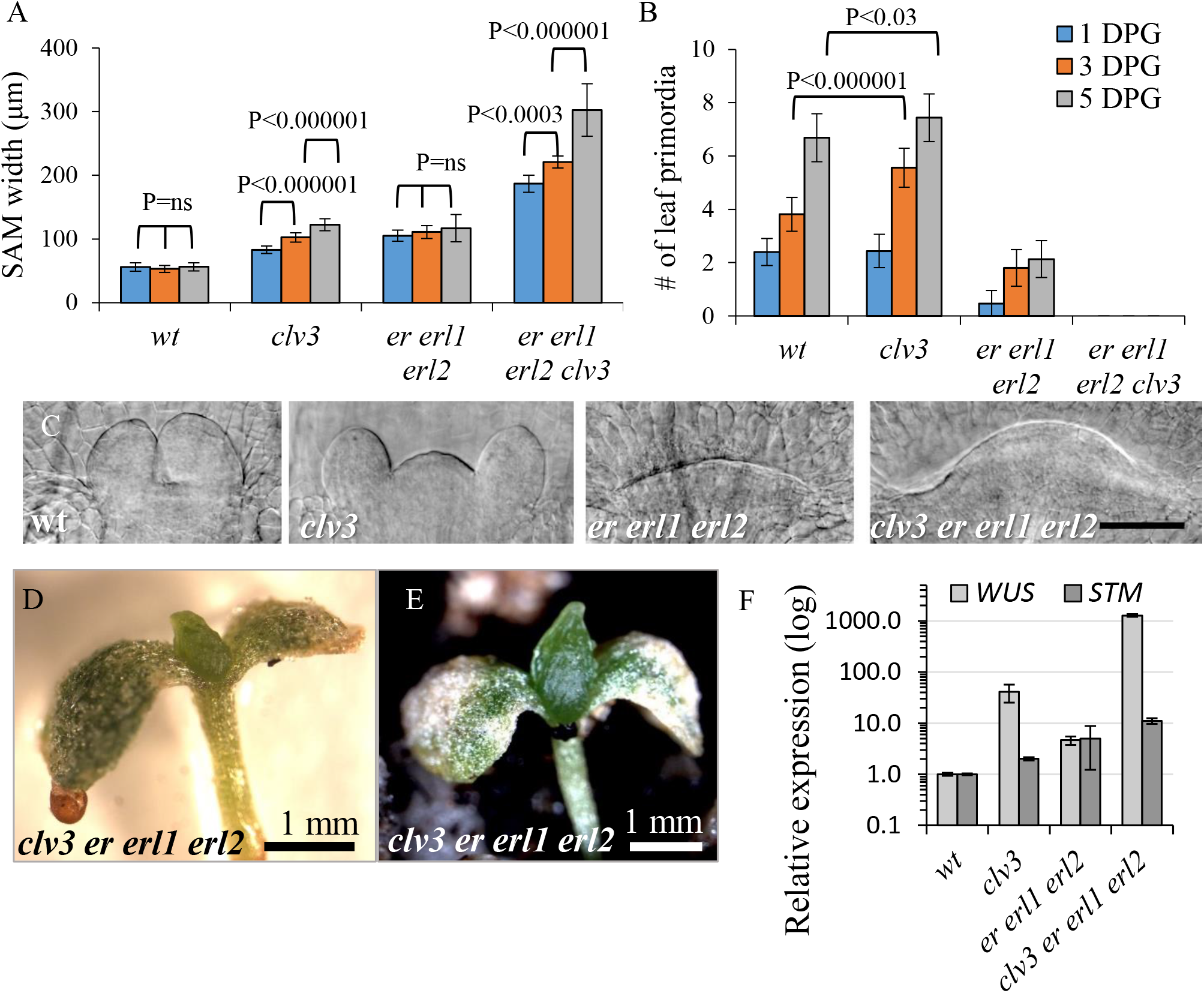
*CLV3* and *ERfs* synergistically regulate SAM size and leaf initiation. (A and B) Comparison of SAM width and leaf initiation. The rate of leaf primordia initiation was determined by DIC microscopy of fixed samples. A primordium was defined as a bulge over 15 μm. Values are mean ± SE, n=6-18, DPG (days post germination). An absence of bars for *er erl1 erl2 clv3* represent a complete absence of leaf primordia in that genotype at that age. The brackets represent results of statistical analysis using Student’s *t*-test. (C) The SAM of dark grown seedlings 1 DPG. All images are at the same magnification. Bar= 50μm. (D) 29 DPG *clv3 er erl1 erl2* plant. (E) 42 DPG *clv3 er erl1 erl2* plants. (F) RT-qPCR analysis of *WUS* and *STM* in above-ground organs of 5-DPG seedlings of wild-type (wt) and mutants as indicated. The average of three biological replicates is presented. Error bars represent SD.

The most dramatic phenotype is observed when both signaling pathways are deactivated. The *clv3 er erl1 erl2* mutants have a considerably larger SAM immediately after germination (Fig. 1A). Post embryonically, the SAM increases in size dramatically overtime but does not form organs (Fig. 1D–E). In rare occasions the mutant will form one or two leaves or produce structures resembling stigmas, but it never forms a stem even after more than 40 days of growth (Fig. S1A and B). The meristematic nature of the dome-like structure in *clv3 er erl1 erl2* is consistent with the presence of cells with dense cytoplasm and without chlorophyll in the outer cell layers (Fig. 3A and Fig.S1C). Moreover, the epidermal layer is composed of very small cells and the guard cells are absent, indicating absence of differentiation (Fig. S1D). Our data are consistent with the larger *clv3 er erl1 erl2* SAM in 10-day-old seedlings described previously (Kimura et al., 2018). The synergistic function of CLV3 and ERfs in the SAM is also evident in the *clv3 er erl2* mutant: while the *er erl2* mutant has a meristem indistinguishable from the wild type, these two mutations enhance the width of the *clv3* SAM, and *er erl2* reduces leaf initiation in the *clv3* background (Fig. S1E and F). Finally, *CLV3* regulates SAM size and leaf initiation in concert with the meristematic ERf ligands *EPFL1*, *EPFL2*, *EPFL4*, and *EPFL6*. The size of the SAM is dramatically increased in *clv3 epfl1 epfl2 epfl4 epfl6* plants, comparable to the increase we observed in *clv3 er erl1 erl2* mutants (Fig. 2). Similar to our previous findings (Kosentka et al., 2019), the four EPFLs function redundantly in regulation of SAM size as we observed a drastic increase only in the pentuple *clv3 epfl1 epfl2 epfl4 epfl6* mutant.

**Fig. 2.**
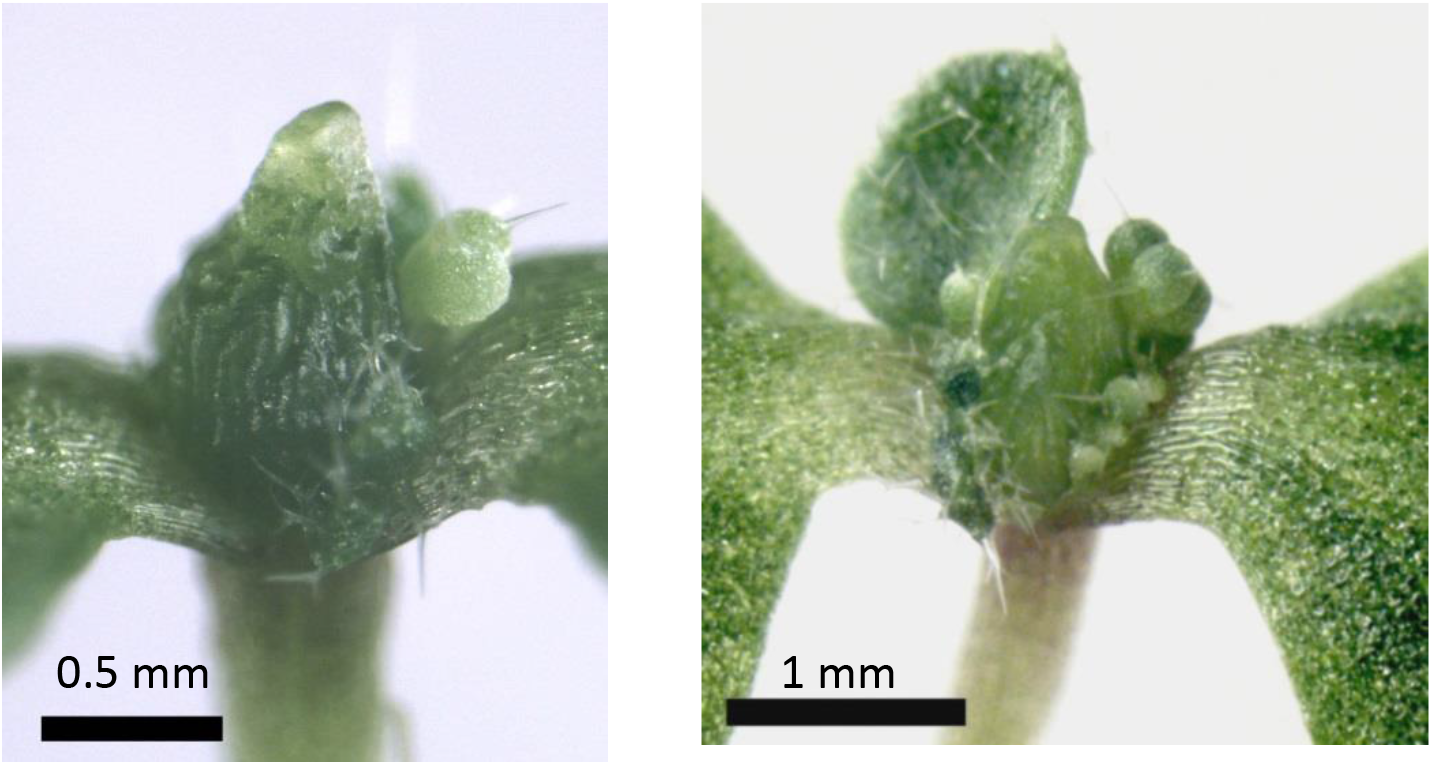
*CLV3*, *EPFL1, EPFL2, EPFL4, and EPFL6* synergistically regulate SAM size. 19 day old *clv3 epfl1 epfl2 epfl4 epfl6* seedlings.

**Fig. 3.**
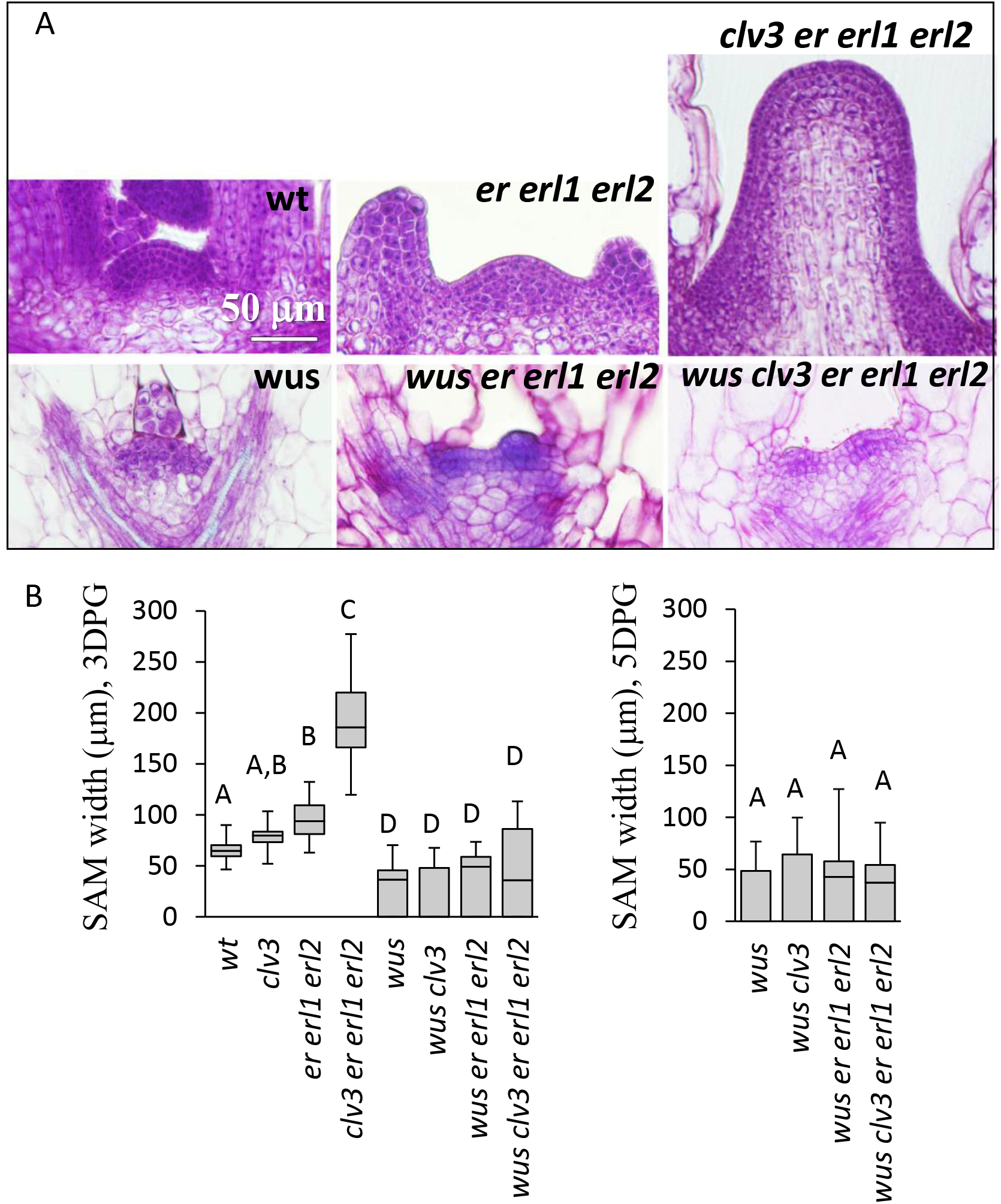
*wus* is epistatic to *er erl1 erl2*. (A) Median sections of shoot apexes of 5 DPG seedlings. All images are at the same magnification. (B) SAM width measurements performed by DIC microscopy using 3 DPG (left) and 5 DPG (right) seedlings. N=15-36. The median is indicated as a thick horizontal line, upper and lower quartiles are represented by the top and the bottom of the boxes, and the vertical lines designate the maximum and the minimum. Different letters indicate significant difference at P < 0.01, as determined by one-way ANOVA with Tukey post-hoc test.

Taken together, these findings indicate that *ERf* and *CLV3* signaling pathways synergistically restrict SAM size. The extent of their individual contributions varies at different developmental stages. Before germination both contribute to SAM size maintenance with ERfs playing the primary role. After germination their roles switch, with CLV3 playing the dominant role and ERf signaling becoming auxiliary.

### The *wus* mutation is epistatic to *er erl1 erl2*

CLV3 is known to regulate meristem size by inhibiting expression of *WUS* (Brand et al., 2000; Muller et al., 2006; Schoof et al., 2000). While expression of *WUS* is increased in an *er erl1 erl2* background, the increase is relatively moderate: 4-6 times more at five DPG (Chen et al., 2013; Uchida et al., 2013) (Fig. 1F). The significance of ERfs for regulation of *WUS* expression becomes more evident in the absence of CLV3 signaling. In 5 DPG *clv3 er erl1 erl2* seedlings we observed up to a ~30x increase in *WUS* expression compared to the single *clv3* mutation and a ~1000x increase over the wild type (Fig. 1F). The entire increase of *WUS* expression in *clv3 er erl1 erl2* is unlikely to be due to a bigger meristem size since the other meristematic marker *STM* increases only ~5.5x compared to *clv3* and 11x compared to the wild type. The large increase of *WUS* expression in *clv3 er erl1 erl2* suggests that its expression is synergistically regulated by CLV3 and ERfs.

To study genetic interactions between *ERfs* and *WUS* and to compare them with *CLV3* and *WUS* genetic interactions, we measured SAM size in *wus*, *wus er erl1 erl2*, *wus clv3*, and *wus clv3 er erl1 erl2* mutants at three and five DPG. While SAM width varied significantly in individual seedlings (Fig. S2A), the four mutants were statistically indistinguishable (Fig. 3B) suggesting that during early seedling growth *wus* is epistatic to both *clv3* and *er erl1 erl2*. This conclusion is supported by histological analysis: the shoot apices of *wus*, *wus er erl1 erl2*, and *wus clv3 er erl1 erl2* mutants did not have the classic dome-like SAM structure consisting of multiple layers of small, evenly shaped, and tightly packed stem cells (Fig. 3A). In the mutants, the shoot apices were composed of only two layers of small cells with some of those cells dividing periclinally, a sign of premature differentiation (Fig. S2B). A previous analysis of *wus er erl1 erl2* using 10 DPG seedlings indicated that its SAM is bigger than that of *wus*, suggesting additive effects of ERf and WUS (Kimura et al., 2018). This conclusion was supported by the ability of *er erl1 erl2* mutations to partially rescue initiation of stamens and carpels in the *wus* background (Kimura et al., 2018). However, our more comprehensive data and analysis of *wus er erl1 erl2* does not support the hypothesis of additive ERf and WUS interactions. At ten DPG in many *wus er erl1 erl2* seedlings we observed a narrow region between forming leaf primordia (Fig. S2C). While in some seedlings the area between forming leaves was indeed enlarged, it did not contain stem cells with the characteristic dense cytoplasm (Fig. S2C). Based on morphology, cells in that region are differentiated: they are highly vacuolated, and some L2 layer cells divide in orientations other than anticlinal. We did occasionally observe meristem-like aggregations of small cells with dense cytoplasm; however, those structures were always small in diameter and asymmetrically localized, often at the axil of a leaf. These structures are either axillary meristems or leaf primordia arising from a few erratically localized stem cells. Thus, while initiation of new meristematic regions or leaf primordia might be altered in *wus er erl1 erl2* compared to *wus*, there is no rescue of the central zone maintenance. Our analysis of *wus er erl1 erl2* flower structure in two-month old plants indicated that ERf family mutations were unable to rescue carpel or stamen initiation in the *wus* background (Table S1). While analyzing flower development we observed formation of stigma-like structures at the tips of sepals and the formation of stigma-like tissue in the area of the SAM in older *wus er erl1 erl2* plants, but in flowers that emerge soon after bolting we never observed the formation of carpels. Although we used the same alleles of *WUS* and *ERfs*, we cannot reproduce the *wus er erl1 erl2* flower structure data described by Kimuta and colleagues (Kimura et al., 2018). In sum, our data indicate that *wus* is epistatic to *er erl1 erl2* in regulation of the SAM central zone width and in the flower meristem.

### ERf signaling inhibits expression of CLV3 and WUS

The expression of both *CLV3* and *WUS* are increased in the *er erl1 erl2* background, with *in situ* hybridization and promoter GUS fusions showing expansion of their expression in the lateral orientation (Chen et al., 2013; Kimura et al., 2018; Uchida et al., 2013). While *epfl1 epfl2 epfl4* and *epfl1 epfl2 epfl6* mutants exhibit only a very slight increase in SAM size (Kosentka et al., 2019), they express *CLV3* at a considerably higher level (Fig. 4A). Is the increase of *WUS* and *CLV3* expression in those mutants a consequence of an enlarged SAM or does ERf signaling inhibit expression of these two genes? To answer this question, we treated *epfl1 epfl2 eplf4* seedlings exogenously with either the EPFL4 peptide or the EPFL6 peptide for 6 hours. RT-qPCR analysis revealed a significantly decreased expression of *WUS* and *CLV3* in response to both peptides (Fig. 4B). Several other genes that have altered expression in *er erl1 erl2* such as *STM* (Fig. 1F), *MONOPTEROS (MP)* (Chen et al., 2013), and *CLV1* (Fig. S3), did not change expression after the peptide treatment (Fig. 4B), suggesting specificity in the downregulation of *CLV3* and *WUS*. The decrease in *WUS* and *CLV3* mRNA levels was dependent on the presence of functional ERf receptors, since it was not observed when *er erl1 erl2* seedlings were treated with EPFL4 (Fig. 4C).

**Fig. 4.**
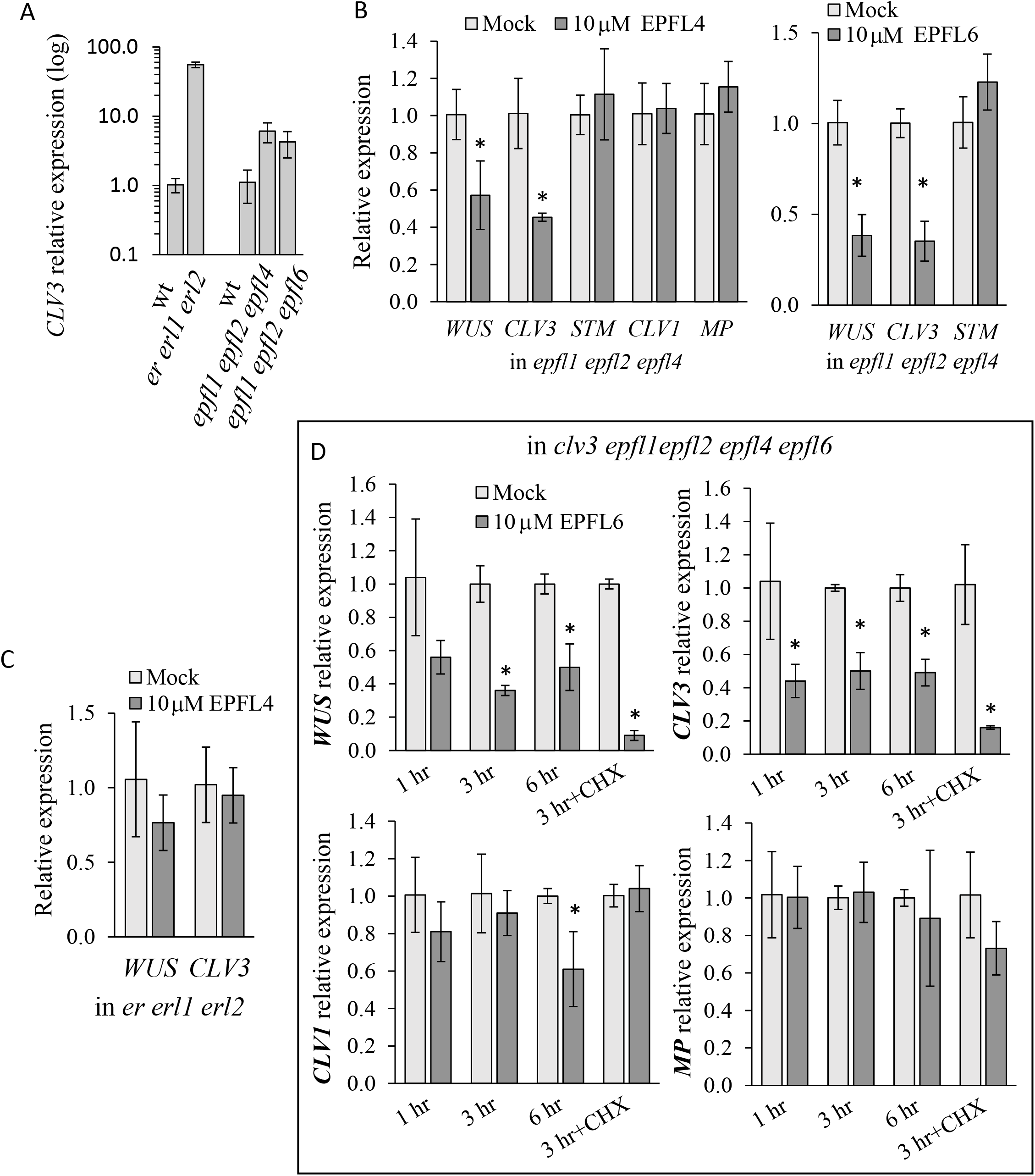
*WUS* and *CLV3* are targets of the ERf signaling pathway. (A) *CLV3* relative expression levels in above-ground parts of 5DPG seedlings in genotypes as indicated (B, C, and D). Relative expression levels of selected mRNAs in above-ground organs of 3 DPG *epfl1 epfl2 epfl4 (B), er erl1 erl2 (C) or clv3 epfl1 epfl2 epfl4 epfl6 (D)* seedlings after treatment with 10μM EPFL4 or EPFL6 peptides as indicated compared to mock treatment. (B and C) Seedlings were treated with peptides for 6 hours. (D) Seedlings were treated with peptide from 1 to 6 hours as indicated. +CHX indicates treatment with 10μM cycloheximide. Data are shown as means ± SD. ACTIN2 was used as an internal control. Values significantly different from mock (P<0.05) as determined by Student *t*-test are indicated by asterisks.

Next, we analyzed whether the ability of EPFLs to suppress expression of *WUS* and *CLV3* is dependent on CLV3 function. The *clv3-9* allele caries a point mutation in the coding region of *CLV3* that disrupts its function but not expression. The *clv3 epfl1 epfl2 epfl4 epfl6* mutant was treated with EPFL6 for one, three, and six hours. The one-hour treatment produced a statistically significant decrease in *CLV3* levels, and the three-hour treatment in the *WUS* levels. Interestingly, the longer treatment did not correlate with a further reduction of *CLV3* or *WUS* expression. At the same time, a treatment of seedlings with EPFL6 in the presence of the translational inhibitor cycloheximide had a very strong impact on the steady-state mRNA levels of *WUS* and *CLV3*. *WUS* and *CLV3* levels decreased approximately eleven and six times, respectively, after three hours of treatment (Fig. 4D). Expression of two other analyzed genes, *MP* and *CLV1*, did not change (Fig 4D). Collectively, our data imply that *WUS* and *CLV3* are downstream targets of the ERf signaling pathway, and the ability of ERfs to inhibit *WUS* and *CLV3* expression is independent of protein biosynthesis.

The *er erl1 erl2* mutant has a reduced sensitivity to CLV3 peptide (Kimura et al., 2018) suggesting that ERfs might have additional roles in regulation of the CLV3 signaling pathway. The reduced sensitivity of the mutant to CLV3 might be related to reduced expression of several CLV3 receptors: CLV1, BAM1, and BAM2 (Fig. S3). However, the role of ERfs in *CLV1* expression is likely to be indirect and complex. While in the *epfl1 epfl2 epfl4* background we did not observe any effects of EPFLs on *CLV1*, in *clv3 epfl1 epfl2 epfl4 epfl6* after six hours of treatment we detected, instead of an increase, a very small decrease in *CLV1* expression (Fig. 4D).

## DISCUSSION

Our analysis of genetic interactions shows that there is a synergy between ERf/EPFL and CLV3 function in the SAM. While *clv3* and *er erl1 erl2* mutants have bigger SAMs, they form stems, leaves, and flowers. In the quadruple mutant the growth of the meristem is unrestricted, one or two leaves form only sporadically, and cells at the periphery of the meristem are unable to differentiate into internode tissues. Synergistic phenotypes most often result from redundancy between paralogs or when pathways converge on a specific node (Pérez-Pérez et al., 2009). Since WUS is the core regulator of SAM size and is the primary target of CLV3, we investigated whether the two signaling pathways converge on that transcription factor. The genetic analysis determined that *wus* is epistatic to *er erl1 erl2* suggesting that WUS could be a downstream target of ERf/EPFL pathway. Treatment of seedlings with exogenous EPFLs for 3 or 6 hours reduced steady-state levels of *WUS* mRNA only in the presence of functional ERf receptors. Considering that the average length of the cell cycle in the SAM is over 30h (R. Jones et al., 2017), the decrease of *WUS* accumulation cannot be attributed solely to a decrease in the size of the SAM. Moreover, when cycloheximide was included in the treatment EPFLs were still able to change steady-state levels of *WUS*, suggesting that EPFLs control *WUS* independently of new protein biosynthesis. Consistent with previously published data (Gordon et al., 2009), we noticed an increased accumulation of *WUS* in the presence of cycloheximide (Fig S3A). Our experiments do not distinguish whether ERf/EPFL signaling controls *WUS* transcription or its mRNA stability. We also do not know why EPFL6 has a stronger impact on *WUS* expression in the presence of cycloheximide. Perhaps there is a negative unstable regulator of *WUS*, and during cycloheximide treatment its concentration drops which enhances the impact of EPFLs on *WUS* transcription. Alternatively, an EPFL-induced increase in *WUS* mRNA degradation might be more evident if an inhibition of translation elongation alters stability of *WUS* mRNA.

It has previously been noticed that ERECTA and CLV3 function along different spatial axes: CLV3 preferentially regulates meristem height and ERECTA regulates meristem width (Mandel et al., 2016). Four ligands that regulate activity of ERfs in the SAM are mostly expressed at the periphery of the meristem and are excluded from the central zone and organizing center (Kosentka et al., 2019). *EPFL1* expression in the peripheral zone under the *KANADI* promoter fully rescues meristematic defects of *epfl1 epfl2 epfl4 epfl6* (26). In contrast, while ERfs are endogenously expressed throughout the SAM, their function in the center of the meristem is critical for SAM maintenance. *ERECTA* expressed under the *CLV3* promoter rescues meristematic defects significantly better compared to its expression under the *KANADI* promoter (Kosentka et al., 2019). The distinct expression of ERfs and EPFLs in the SAM is similar to their distinct expression during leaf tooth initiation, where ERfs are expressed more strongly at the tip of the tooth while EPFL2 is excluded from the tip and is expressed in the surrounding sinus tissues (Tameshige et al., 2016). Expression of *ERECTA* under the *DR5* promoter exclusively at the tip of the leaf tooth rescues its growth (Tameshige et al., 2016), just as expression of *ERECTA* under the *CLV3* promoter in the center of the meristem rescues meristematic phenotypes (Kosentka et al., 2019). Previously we proposed that the signaling occurs in the peripheral region of the SAM where expression of EPFL and ERf overlaps, a pattern similar to that observed in the leaf tooth.

We propose that ERfs restrict the size of the central zone by inhibiting expression of *CLV3* and *WUS* in the periphery of the SAM (Fig. 5). This control is especially important during establishment of the SAM during embryogenesis. After germination CLV3 signaling can partially substitute for ERfs in the lateral inhibition of *WUS* expression. However, when both signaling pathways are disrupted, as in the *clv3 er erl1 erl2* mutant, *WUS* expression becomes rampant, and the SAM expands without restraint. Increased expression of *CLV3* in the L1 layer of *er erl1erl2* and *wus er erl1 erl2* (Kimura et al., 2018) suggests that an as-yet unidentified signal induces *CLV3* expression in the epidermis, consistent with a previously proposed model (Gruel et al., 2016). The functional significance of ERfs in suppressing *CLV3* expression in the periphery of the meristem is yet to be determined. A computational model might help to address this question in the future.

**Fig. 5.**
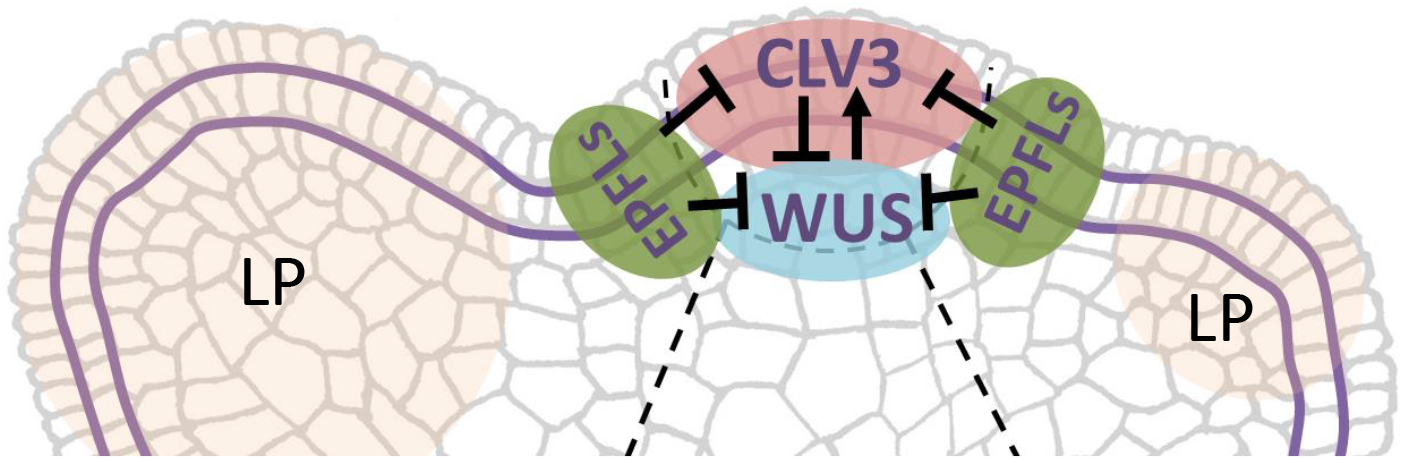
Model for the role of ERf signaling in regulation of shoot meristem maintenance. EPFL signals originating from the periphery of the SAM activate the ERf signaling cascade that inhibits expression of *CLV3* and *WUS* thus restricting the width of the central zone. LP – leaf primordium.

## Materials and Methods

### Plant Materials and Growth Conditions

The Arabidopsis (Arabidopsis thaliana) ecotype Columbia was used as the wild type. The following mutants used in the study have been described previously: *er-105 erl1-2 erl2-1*(Shpak et al., 2004), *epfl1 epfl2 epfl4*, *epfl1 epfl2 epfl6* and *epfl1 epfl2 epfl4 epfl6* (Kosentka et al., 2019), *clv3-9* (Nimchuk et al., 2015), and *wus* null allele (SAIL_150_G06) (Sonoda et al., 2007). They are all in the Columbia background.

To create *clv3 er erl2, clv3 er erl1 erl2*, and *wus er erl1 erl2* plants *clv3-9* and *wus*/+ were crossed with *er erl1*/+ *erl2*. To create *wus clv3* plants *clv3-9* was crossed with *wus*/+. To create *wus clv3 er erl1 erl2* plants *clv3 er erl1*/+ *erl2* was crossed with *wus*/+ *er erl1*/+ *erl2*. The higher order mutants were identified in subsequent generations based on the phenotype. The homozygous status of *wus* and *erl1* were confirmed when necessary by genotyping as described previously (Sonoda et al., 2007), (Kosentka et al., 2017). To create *clv3 epfl1 epfl2 epfl4 epfl6* plants the *clv3-9* mutant was crossed with *epfl1*/+ *epfl2 epfl4 epfl6*. The *epfl* mutations were genotyped as described previously (Kosentka et al., 2019).

Plants were grown as described elsewhere (Kosentka et al., 2017) under an 18-h light/ 6-h-dark cycle (long days) at 21°C. For analysis of SAM size and leaf initiation seedlings were grown on modified Murashige and Skoog medium plates supplemented with 1% (w/v) sucrose. For all experiments, seeds were stratified for 2 days at 4°C before germination.

To analyze expression of genes after EPFL4 and EPFL6 treatment, *epfl1 epfl2 epfl4* mutants were grown on modified Murashige and Skoog medium plates for 5 days (3DPG). Then 60 seedlings (EPFL4 treatment) or 10 seedlings (EPFL6 treatment) per biological replicate were transferred to 1 ml of liquid Murashige and Skoog medium containing 10μM of EPFL4 or EPFL6. The purification of EPFL4 and EPFL6 peptides has been described previously (Lin et al., 2017). EPFL peptides were dissolved in 10 mM Bis-Tris, 100 mM NaCl, pH=6.0. For mock treatment, a buffer solution of equal volume was added to the medium (92.6 μl in the EPFL4 experiment and 8.7 μl in the EPFL6 experiment). Plants were pretreated with 10 μM cycloheximide with and without 10μM of EPFL6. For each treatment there were 3 biological replicas.

### Microscopy

To measure leaf initiation and SAM size, one, three, and five DPG seedlings were fixed overnight with ethanol: acetic acid (9:1 [v/v]). After fixation, samples were rehydrated with an ethanol series to 30% (v/v) ethanol and cleared in chloral hydrate solution. Chloral hydrate: water: glycerol 8:1:1 [w/v/v] solution contained KOH at 10mM concentration to prevent degradation of tissues due to high acidity of chloral hydrate. In our experience the acidity of chloral hydrate (Sigma-Aldrich) varies from batch to batch, and the necessity to add KOH should be tested experimentally. Microscopic observations of meristematic regions by DIC microscopy were performed as described previously (Chen et al., 2013). Tissue samples were fixed overnight in acetic acid: ethanol (1:9) at RT, dehydrated with a graded series of ethanol, and infiltrated with polymethacryl resin Technovit 7100 (Heraeus Kulzer, Wehrheim, Germany) followed by embedding and polymerization in Technovit 7100. Seven-micrometer sections were prepared using a Leica RM-6145 microtome (Wetzlar, Germany). The tissue sections were stained with 0.02% toluidine blue O and observed under bright-field illumination. Pictures of older seedlings and the analysis of flower structure was done using a Leica MZ16 FA stereomicroscope.

### Quantitative RT-PCR Analysis

Total RNA was isolated from the aboveground tissues of seedlings using the Spectrum Plant RNA Isolation Kit (Sigma-Aldrich). The RNA was treated with RNase-free RQ1 DNase (Promega). First-strand complementary cDNA was synthesized with LunaScript™ RT SuperMix Kits (New England Biolabs). Quantitative PCR was performed with a CFX96 Touch Real-Time PCR Detection System (Bio-Rad) using Sso Fast EvaGreen Supermix (Bio-Rad). Each experiment contained three technical replicates of three biological replicates and was performed in a total volume of 10 μL with 4 μL of 10x or 50x diluted cDNA. Cycling conditions were as follows: 3 min at 95°C; then 40 repeats of 10 s at 95°C, 10 s at 52°C for *ACTIN2* and *STM*; 10 s at 55°C for *WUS*; 10s at 57°C for *MP*; 10 s at 50°C for *CLV1*, *BAM1*, *BAM2*, and *BAM3*, and 10 s at 68°C, followed by the melt-curve analysis. Cycling conditions for *CLV3* were 3 min at 95°C; then 40 cycles of 95°C for 10 s, and 60°C for 10 s, followed by the melt-curve analysis. Primers for *ACTIN2*, *STM*, *WUS*, and *MP* (Chen et al., 2013) as well as for *CLV3* (Chiu et al., 2007) have been described previously. Primers for *CLV1*, *BAM1*, *BAM2*, and *BAM3* were as in (Nimchuk et al., 2015). They are CLV1qPCRf GGTTCAATCCCTACCGGAAT and CLV1qPCRr CCAAGAATTGACCACCGAGT; BAM1qPCRf TCTCCGGTCCATTAACTTGG and BAM1qPCRr CGAAACTCGCTGGAATCTCT; BAM2qPCRf TCAATGGGTGAGAAGCATGA and BAM2qPCRr CAGAGCAACGCAACGTAGAA; BAM3qPRCf CGCTTACGACAACAGCTTCA and BAM3qPRCr GGGATCTCACCGTCGAAGTA. The fold difference in gene expression was calculated using relative quantification by the 2^−ΔΔCT^ algorithm.

## Acknowledgments

We thank Zachary Nimchuk for sharing with us seeds of *clv3-9* and *wus* (SAIL_150_G06). We thank Richard Maradiaga and Timothy Herman for technical assistance.

## Author contributions

E.S. conceived and directed the project; E.S and C.J. designed experiments; L.Z., D.D, G.L. and E.S. performed experiments and analyzed data, E.S. and L.Z wrote the paper.

## Data and materials availability

Arabidopsis Genome Initiative numbers for the genes discussed are as follows: BAM1 (AT5G65700), BAM2 (AT3G49670), CLV1 (At1g75820), CLV3 (At2g27250), ER (At2g26330), ERL1 (At5g62230), ERL2 (At5g07180), EPFL1 (At5g10310), EPFL2 (At4g37810), EPFL4 (At4g14723), EPFL6 (At2g30370), and WUS (At2g17950). All data are available in the main text or the supplementary materials.

## This article contains supporting information

Figs. S1 to S3

Table S1

